# Distinct Repeat Architecture Landscapes in the Proteomes of Protozoan Parasites

**DOI:** 10.64898/2026.01.20.700692

**Authors:** Hirotaka Matsumoto, Jing Hong

## Abstract

Protozoan parasites cause major infectious diseases and pose persistent challenges to global health, particularly the emergence of drug-resistant strains. Tandem repeats (TRs) and other repetitive architectures are widespread in proteomes, especially in protozoan proteins, where they have been implicated in host–parasite interactions, immune evasion, and antigenicity. However, repeat-containing proteins (RPs) exhibit highly diverse architectures that often extend beyond the simple reiteration of a single motif, making comprehensive and quantitative characterization challenging.

In this study, we performed bioinformatics analysis of repeat architectures in protozoan proteins. In addition to the established repeat-detection approaches, we developed a new algorithm, Drepper, which quantifies repeat-architecture complexity. By integrating diverse repeat-related features, we clustered RPs across species and identified distinct groups associated with parasite lineages. Notably, we detected a *Plasmodium*-specific RP cluster and a *Trypanosoma*/*Leishmania*-specific RP cluster; both were characterized by large repeat regions but exhibited contrasting repeat-structure complexity. The *Plasmodium*-specific RPs showed high complexity, whereas the *Trypanosoma*/*Leishmania*-specific RPs displayed significantly low complexity. Functional enrichment analyses indicated that these lineage-associated clusters were enriched in parasite-specific factors. Furthermore, evolutionary analyses suggested that low-complexity repeat architectures may be actively maintained through concerted evolution.

Taken together, our results reveal lineage-specific strategies in protozoan repeat architectures and provide a quantitative framework for studying their biological and evolutionary roles.

## Introduction

Protozoan parasites are parasitic unicellular eukaryotes including malarial parasites and *Leishmania*, many of which cause severe infectious diseases, including malaria caused by *Plasmodium* and leishmaniasis caused by *Leishmania*. These diseases result in a substantial number of deaths and are a major threat to global public health. Malaria alone affects hundreds of millions of individuals worldwide, with approximately 600,000 deaths annually (1). In addition, an estimated 50,000–90,000 new cases of visceral leishmaniasis occur worldwide annually, with fatalities in over 95% of the cases when untreated (2). In recent years, the emergence and spread of artemisinin-resistant malaria parasites have been reported, posing a new challenge for malaria control. The development of novel antiparasitic drugs and vaccines is urgently required to combat protozoan infectious diseases, including the expansion of drug-resistant strains. To achieve this, we require a detailed understanding of how protozoa invade host cells and evade or modulate host immune responses, at the molecular and cellular levels.

Tandem repeats (TRs) are sequence architectures in which a specific amino acid sub-sequence, referred to as a repeat unit, is repeated iteratively. Proteins that contain TRs or other repetitive sequence architectures are collectively referred to as repeat-containing proteins (RPs). Early estimates suggested that approximately 14% of proteins, universally, are RPs (3), whereas more recent studies suggest that a substantially larger proportion of proteins contain repeat architectures (4). Such repeat architectures form domains that mediate interactions with other proteins or nucleic acids (5). Although repeat regions often lack stable tertiary structures and tend to be intrinsically disordered (6), they can function as scaffolds for molecular recognition and signal transduction (7). Based on these properties, repeat architectures have been actively exploited in the design of artificial proteins (8). Several RPs have been reported in protozoa, and many proteins in malaria parasites are composed of repetitive low-complexity sequences (9). For instance, the circumsporozoite protein (CSP), a major target of malaria vaccines, contains prominent repeat regions. Notably, the repeat sequences of CSP differ between *Plasmodium falciparum* and *Plasmodium vivax*, suggesting that a detailed understanding of repeat features is important for vaccine development (9). In addition, repeat-containing proteins in protozoa are thought to play critical roles in processes such as cell adhesion, host cell invasion, and immune evasion (10). Repeat features differ between intracellular parasites, extracellular parasites, and free-living protists, suggesting a link between repeat architectures and parasitic lifestyles. Moreover, TR-containing proteins are frequent targets of B-cell immune responses (11, 12), and high antigenicity of TR-containing proteins has been reported in *Leishmania* and *Trypanosoma* species (13, 14). Although the numbers of RPs in *Leishmania* and *Trypanosoma* are lower than that in malaria parasites, these organisms often include RPs with exceptionally large TR regions. This observation highlights species-specific differences in repeat architectures and suggests that such differences may be linked to distinct parasitic strategies (14).

Repeat sequences are prone to rapid changes through expansion, contraction, and mutation, and are therefore considered potential drivers of evolutionary innovation (15, 16). In protozoa, increasing attention is being paid to the evolution of repeat sequences and their effects on parasitism, including species-specific differences in CSP repeats (9, 17–19). In particular, repeat regions undergo concerted evolution, in which repeat units evolve cooperatively rather than independently. This mode of evolution may have a profound effect on host–parasite co-evolution (17).

Taken together, these findings indicate that RPs play an important role in protozoan parasitic strategies. However, the properties and functions of these RPs are highly diverse, and the biological significance of the repeat architectures in protozoa remains unclear.

Bioinformatics tools are essential for detecting and analyzing repeat architectures in protozoan amino acid sequences.

To date, various algorithms have been developed to detect the TR regions, primarily based on sequence-processing approaches (20–23). However, repeat architectures are often more complex than the simple reiterations of a single motif and may exhibit hierarchical or composite structures. Consequently, repeats cannot be treated as a uniform class and require multifaceted characterization and classification.

In this study, we aimed to elucidate the diversity and characteristic features of repeats in protozoan proteins by quantitatively characterizing the repeat architectures from multiple perspectives and performing comprehensive analyses (Fig. 1). In addition to utilizing existing bioinformatics tools, we developed a novel algorithm, Drepper, based on self dot plots, which enables quantitative assessment of repeat architecture complexity. Using a diverse set of repeat-related features, we clustered RPs and identified a *Plasmodium*-specific cluster and a *Trypanosoma*/*Leishmania*-specific cluster. Although most RPs in these clusters contained large repeat regions, *Trypanosoma*/*Leishmania*-specific RPs were characterized by markedly low repeat-structure complexity than RPs in the *Plasmodium*-specific cluster. This work represents the first comprehensive, quantitative comparison of repeat architecture complexity across protozoan lineages, uncovering lineage-specific repeat patterns that are difficult to capture using conventional repeat detection approaches.

**Fig. 1.**
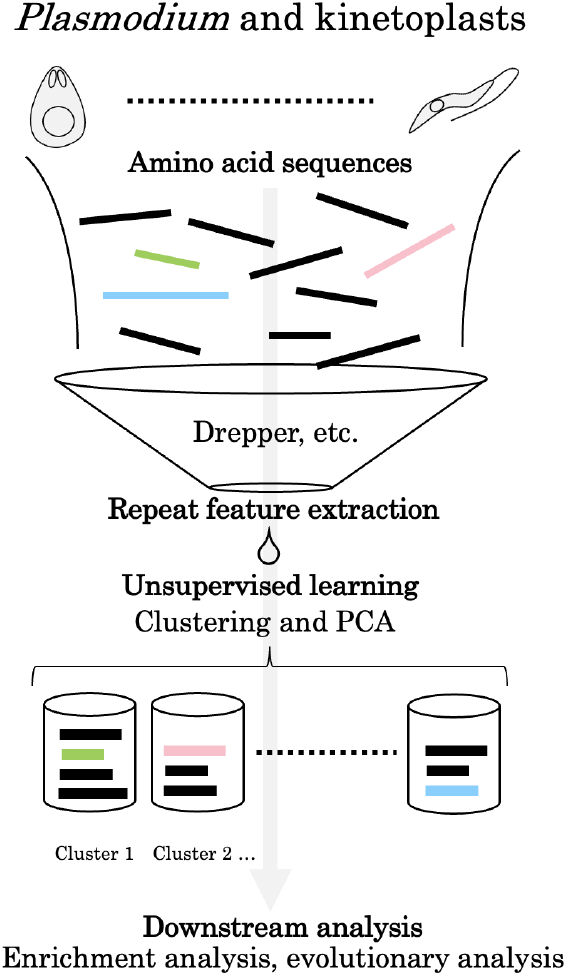
A graphical abstract illustrating the analytical workflow of this study.

## Results

### Overview of repeat architecture analysis

We analyzed amino acid sequences from 12 species of malaria parasites and kinetoplastids (Table 1), quantitatively characterizing repeat architectures using seven repeat-related features derived from existing tools and a newly developed algorithm, as summarized below (details are provided in the Methods section).

**Table 1.**
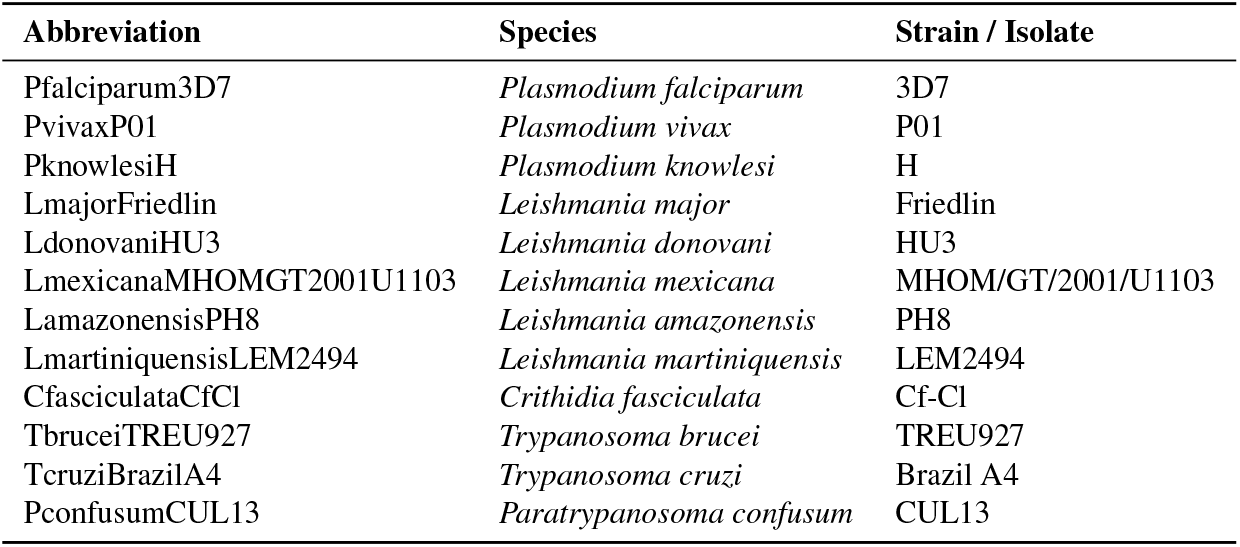
Species and strain information.

| Abbreviation | Species | Strain / Isolate |
| --- | --- | --- |
| Pfalciparum3D7 | *Plasmodium falciparum* | 3D7 |
| PvivaxP01 | *Plasmodium vivax* | P01 |
| PknowlesiH | *Plasmodium knowlesi* | H |
| LmajorFriedlin | *Leishmania major* | Friedlin |
| LdonovaniHU3 | *Leishmania donovani* | HU3 |
| LmexicanaMHOMGT2001U1103 | *Leishmania mexicana* | MHOM/GT/2001/U1103 |
| LamazonensisPH8 | *Leishmania amazonensis* | PH8 |
| LmartiniquensisLEM2494 | *Leishmania martiniquensis* | LEM2494 |
| CfasciculataCfCI | *Crithidia fasciculata* | Cf-CI |
| TbruceiTREU927 | *Trypanosoma brucei* | TREU927 |
| TcruziBrazilA4 | *Trypanosoma cruzi* | Brazil A4 |
| PconfusumCUL13 | *Paratrypanosoma confusum* | CUL13 |

**Length** Amino acid sequence length

**Trcount** Number of TRs

**MaxTRS** Maximum TR region size

**RUP** Purity of the repeat unit sequence

**MaxRUS** Maximum repeat unit size

**Complexity** Complexity of the repeat structure

**TotalRS** Total repetitive region size

### Clustering of RPs

Seven repeat-related features were computed for each RP, and the RPs were clustered based on these features. Fig. 2A shows a visualization of the clustering results projected onto the principal component analysis (PCA) space. Hereafter, Clusters 1 through 7 are denoted as C1–C7, respectively.

**Fig. 2.**
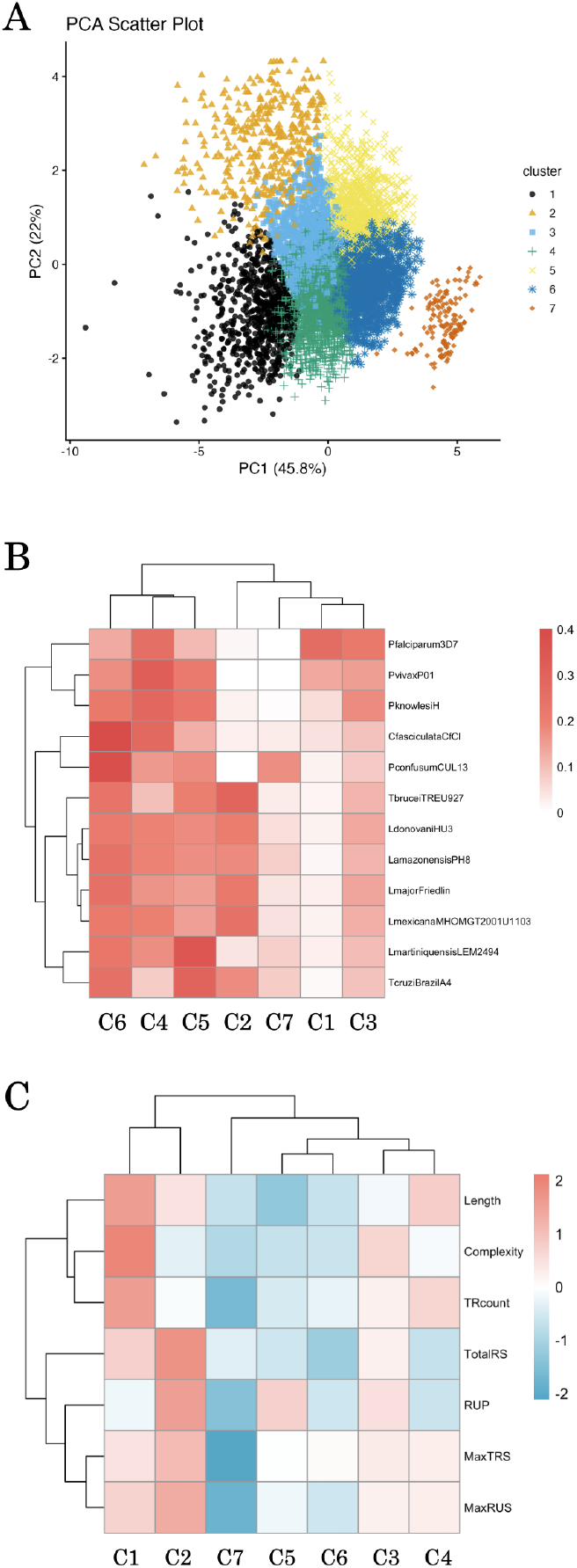
(A) Visualization of amino acid sequence clustering based on repeat-related features projected onto the PCA space. (B) Heatmap showing the species composition of each cluster. (C) Heatmap of standardized mean values of repeat-related features for each cluster.

First, we examined the species composition of each cluster and visualized the results using a heatmap (Fig. 2B). As shown in Fig. 2B, C1 was predominantly composed of *Plasmodium* species, particularly *P. falciparum* (Pfalciparum3D7). In contrast, C2 was enriched in *Trypanosoma* species (TbruceiTREU927 and TcruziBrazilA4) and multiple *Leishmania* species (LmajorFriedlin, LdonovaniHU3, LamazonensisPH8, and LmexicanaMHOMGT2001U1103). The degree of enrichment was statistically assessed using a generalized binomial model (see Methods). In C1, the proportions of Pfalciparum3D7, PvivaxP01, and PknowlesiH were significantly higher than those of other species (*p*< 2 ×10^−16^). Similarly, in C2, the enrichment of *Trypanosoma* and *eishmania* species was highly significant (*p*< 2 × 10^*−*16^).

To examine species-specific trends independent of clustering, we visualized the density distribution of RPs for each species in the PCA space. Representative examples are presented in Fig. 3A (all species are shown in the Supplementary Note). For Pfalciparum3D7, a relatively high density of data points was observed in the regions corresponding to C1, whereas for LmajorFriedlin, a higher density was observed in the regions corresponding to C2.

**Fig. 3.**
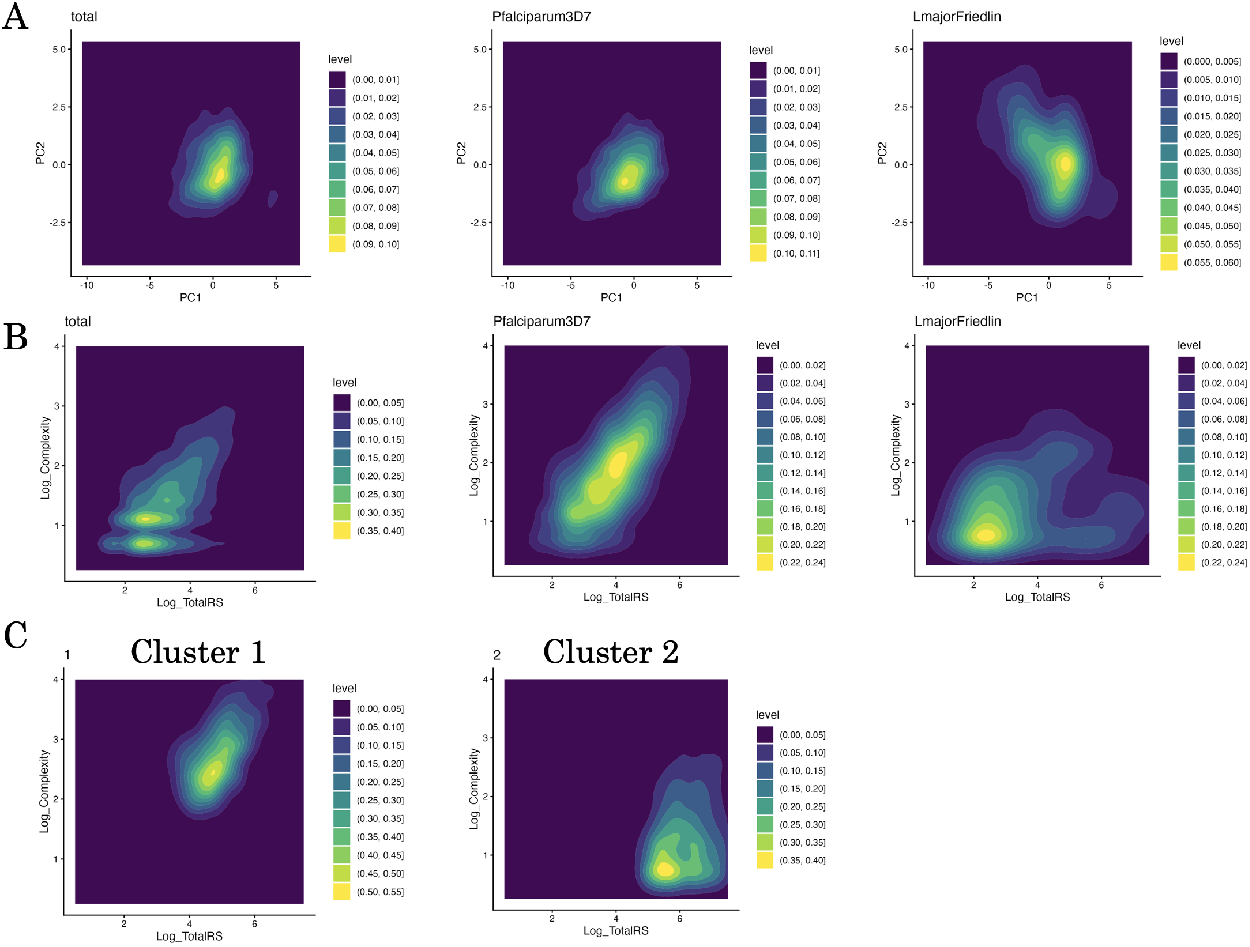
(A) Density distributions of RPs in the PCA space, shown for all data (left), Pfalciparum3D7 only (middle), and LmajorFriedlin only (right). (B) Density plots of RPs with logarithm of TotalRS on x-axis and logarithm of Complexity on y-axis, shown for all data (left), Pfalciparum3D7 only (middle), and LmajorFriedlin only (right). (C) Density plots corresponding to panel B, shown separately for C1 (left) and C2 (right).

To characterize the repeat-related features within each cluster, we calculated the mean values of each feature for each cluster and visualized them using a heatmap (Fig. 2C). Both C1 and C2 exhibited large values of TotalRS, indicating that RPs in these clusters generally contained extensive repeat regions. However, these two clusters showed contrasting feature profiles: C1 was characterized by high complexity and low RUP, whereas C2 showed the opposite trend.

C3 and C4 also exhibited relatively large values of TRcount, MaxTRS, and MaxRUS, although to a lesser extent than C1. Among these, C3 showed high RUP and complexity, whereas C4 showed lower values for these features. C5 and C6 were characterized by smaller TotalRS values, with C5 exhibiting higher RUP and C6 exhibiting lower RUP. C7 exhibited uniformly small values for MaxTRS and related features. As discussed below, repeats in C7 were predominantly dispersed rather than TRs, forming a distinct group that was separated from the other clusters in the PCA space.

To further investigate the relationship between TotalRS and Complexity, we visualized these features for all RPs, and separately for Pfalciparum3D7 and LmajorFriedlin (Fig. 3B. The results for other species are presented in the Supplementary Note). We also visualized the distributions of C1 and C2 (Fig. 3C). In Pfalciparum3D7, complexity tended to increase with increasing TotalRS. In contrast, LmajorFriedlin exhibited a substantial number of RPs with a large TotalRS but low complexity. Accordingly, the sequences in C1 and C2 were clearly separated into groups with high and low complexity at high TotalRS values.Representative examples of RPs and their repeat architectures in C1 and C2 are shown as dot plots in Fig. 4 (examples for the other clusters are provided in the Supplementary Note). Numerous off-diagonal elements were observed in both clusters, indicating the presence of extensive repeat regions. In C1, many RPs contained multiple distinct repeats within a single sequence or exhibited highly complex repeat architectures. In contrast, RPs in C2 frequently formed large repeat regions that were close to perfect repeats.

**Fig. 4.**
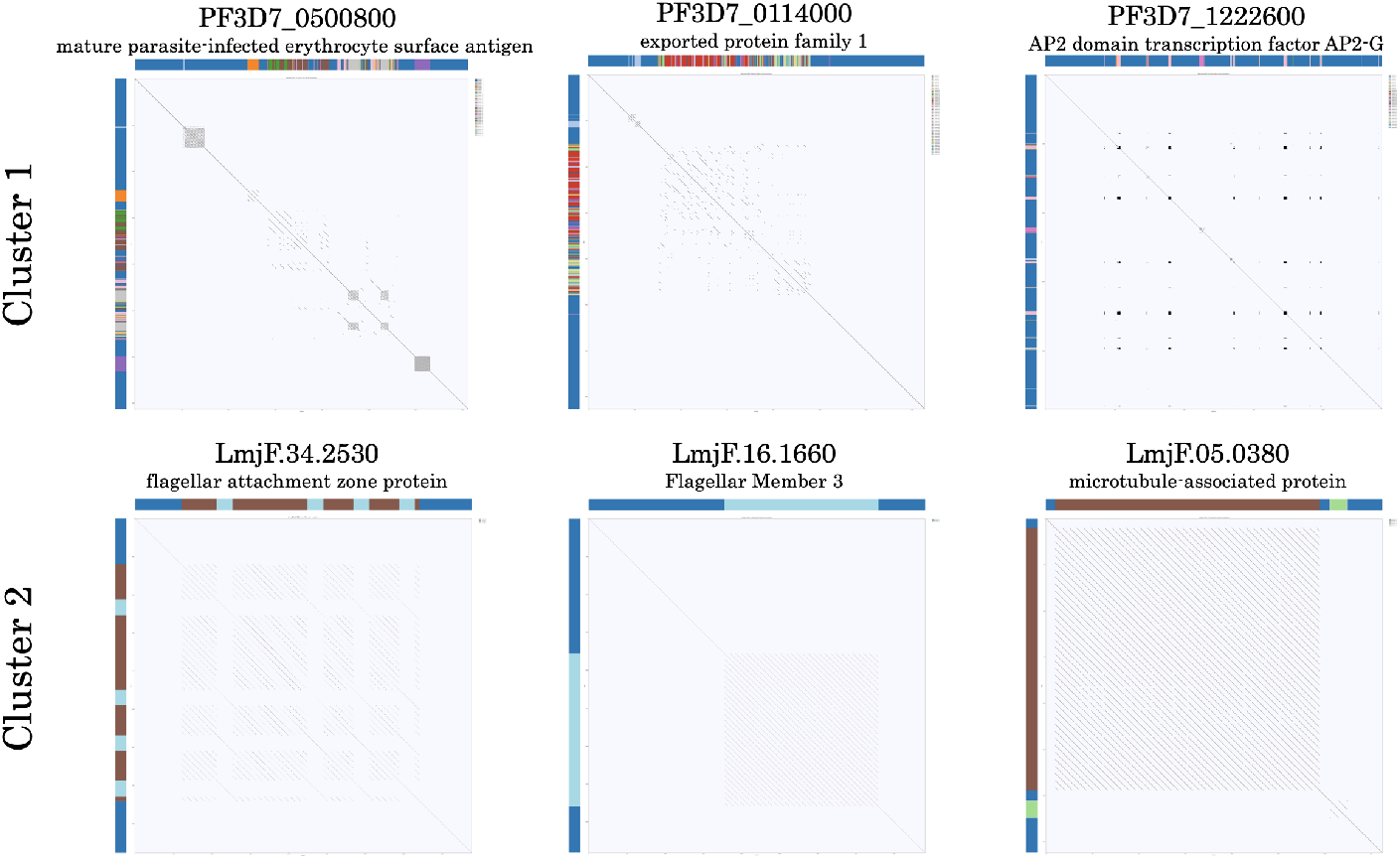
Representative dot plots of RPs in C1 (top) and C2 (bottom). The annotations displayed along the top and left margins of the dot plot, together with their colors, represent the clustering results obtained using Drepper (see Methods section).

Finally, RPs in C7 generally contained repeats that were spatially dispersed along the sequence, rather than forming contiguous TRs (Supplementary Note). Consequently, their repeat architectures differed substantially from the patterns typically detected using tandem repeat detection tools, and these sequences formed a separate distant cluster in the PCA space.

### Functional enrichment analyses

We analyzed the functional characteristics of RPs contained in each cluster based on Gene Ontology (GO) and protein signatures.

GO enrichment analysis was performed for each cluster using two complementary control sets, and the most significantly enriched terms were visualized (Fig. 5; see Methods for details). For C1, GO terms, such as nucleus (GO:0005634) and sequence-specific DNA binding (GO:0043565), were highly enriched. In C2, enriched GO terms included ciliary basal body (GO:0036064), microtubule binding (GO:0008017), and cilium (GO:0005929). Other clusters exhibited distinct enrichment patterns: C3 was enriched for host cell (GO:0043657), C4 for cell adhesion molecule binding (GO:0050839), host cell surface receptor binding (GO:0046789), and infected host cell surface knob (GO:0020030), whereas C7 was enriched for ATPase-coupled transmembrane transporter activity (GO:0042626).

**Fig. 5.**
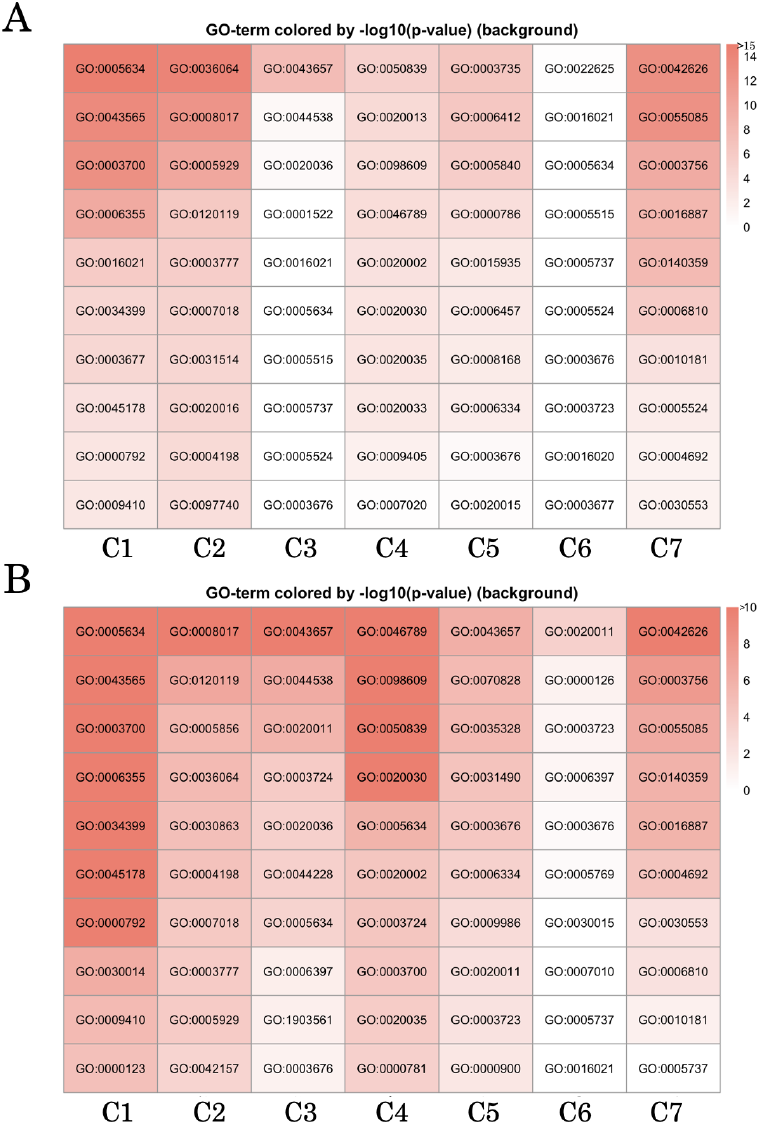
Top 10 Gene Ontology terms enriched in each cluster. Colors represent *p*-values. The top panel (A) presents the results of enrichment analysis using all other clusters as controls, whereas the bottom panel (B) shows that using all genes outside the target cluster, including non-RPs, as controls.

We also performed protein signature enrichment analysis for each cluster using two complementary control sets, and the most significantly enriched terms were visualized (Fig. 6; see Methods for details). In C1, the enriched signatures included the CATH superfamily G3DSA:1.20.5.2050, which contains transcription factors with AP2 domains, as well as the AP2 domain itself (PF00847). In C2, signatures, such as flagellum attachment zone protein 1 (FAZ1; PF23398) and the domain of unknown function DUF7623 (PF24610), were highly enriched. In addition, Mucin-like glycoprotein (PF01456) frequently appeared among the most enriched signatures in C5 and C6. When enrichment was assessed using controls that included non-RPs, the protein signature mobile-lite, which represents intrinsically disordered proteins (IDPs), was consistently enriched across C1 to C6.

**Fig. 6.**
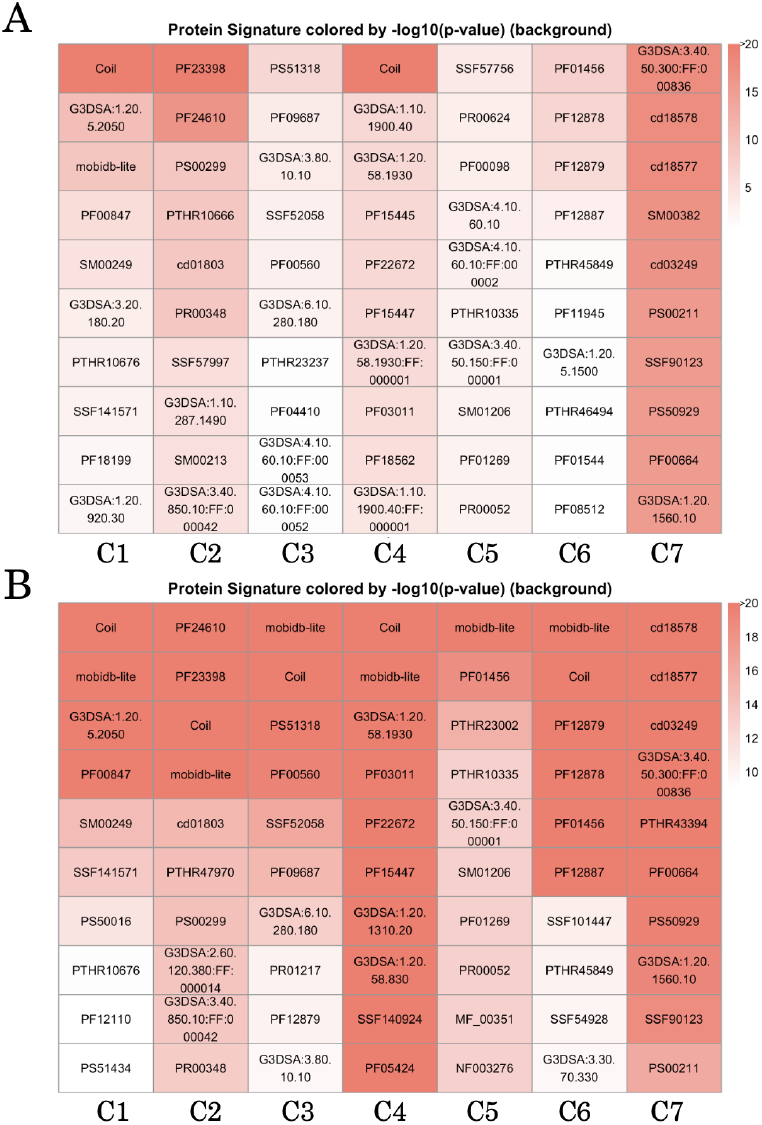
Top 10 enriched protein signatures in each cluster. Colors represent *p*-values. The top panel (A) shows the results of enrichment analysis using all other clusters as controls, whereas the bottom panel (B) presents that using all genes outside the target cluster, including non-RPs, as controls.

C1, which was enriched for *Plasmodium* proteins, showed enrichment of GO terms related to nucleic acid binding and sequence-specific DNA binding; protein signature analysis highlighted domains related to AP2 transcription factors. In *Plasmodium*, the Apicomplexan AP2 (ApiAP2) protein family constitutes the core of sequence-specific transcriptional regulators (24). Our results suggest that these proteins frequently contain large repeat regions with high complexity.

C2, which was enriched for *Trypanosoma* and *Leishmania*, showed consistent enrichment of flagellum-related functions in both GO and protein signature analyses. This suggests that the proteins associated with these functions tend to contain large repeat regions with low complexity, which may contribute to their biological roles. The DUF7623 domain, which is largely specific to Trypanosomatidae and has been reported to be associated with calpain family cysteine proteases (PF00648), is thought to be involved in parasite life-cycle regulation and parasitism, although its precise function remains unknown.

In C3, host cell appeared to be the most enriched GO term. All genes in this category were derived from *Plasmodium* species, including 11 genes from Pfalciparum3D7, 21 from PvivaxP01, and 18 from PknowlesiH, with a total of 50 genes. Among these, 49 were annotated as *Plasmodium*-exported proteins, many of which were associated with the PHIST (Plasmodium helical interspersed subTelomeric) family or classified as hyp1, hyp2, or hyp11 proteins. The remaining gene encodes an MSP7-like protein (PF3D7_1334300).

In C4, the enrichment of terms, such as host cell surface receptor binding (GO:0046789), was largely attributable to the presence of 24 *P. falciparum* PfEMP1 genes in this cluster. In C5 and C6, enrichment of mucin-like glycoprotein (PF01456), a *Trypanosoma*-derived protein family structurally similar to vertebrate mucins, was observed. These genes encode the core proteins of the parasite mucins and may be involved in interactions with mammalian host cells (25).

Finally, when clusters were compared with controls, including non-RPs, enrichment of the mobile-lite protein signature was consistently observed. This finding corroborates previous observations that RPs tend to exhibit intrinsically disordered properties.

### Evolutionary analysis

To investigate the evolutionary characteristics of low-complexity repeat architectures, we performed a comparative analysis of C2 repeat-containing proteins across multiple *Leishmania* species. By comparing orthologous repeat regions among *Leishmania* species enriched in C2, we assessed patterns of repeat unit similarity and divergence within low-complexity tandem repeat regions. Twelve orthogroups were identified, in which orthologs from all four *Leishmania* species were assigned to C2. Among these, we focused on three representative orthogroups: Orthogroup 1 containing LmjF.34.2530 (Flagellar attachment zone protein), Orthogroup 2 containing LmjF.16.1660 (Flagellar Member 3), and Orthogroup 3 containing LmjF.33.3070 (Flagellar Member 8).

First, from the repeat motifs identified in LmajorFriedlin, we selected motifs for which similar repeats were also present in the other three species and enumerated all sub-sequences within the orthologous proteins that were similar to these motifs. These sequence hits were then subjected to multiple sequence alignments, and sequence logos were generated for each species (Orthogroup 1, shown in Fig. 7; results for Orthogroup 2 and 3 are provided in the Supplementary Note). In Orthogroup 1, for example, the first amino acid position was almost uniformly conserved as glutamic acid (E) across all four species, whereas the seventh position differed among the species: E in LmajorFriedlin, K in LdonovaniHU3, and L in both LmexicanaMHOMGT2001U1103 and LamazonensisPH8 (Fig. 7). These results indicate that repeat unit sequences within orthologs exhibit species-specific differences. Based on the multiple sequence alignment results, phylogenetic networks of repeat unit sequences were constructed for each orthogroup (Fig. 8). In all three orthogroups, the repeat unit sequences were clustered predominantly, by species. Moreover, in terms of sequence similarity, LamazonensisPH8 and LmexicanaMHOMGT2001U1103 were relatively closer to each other than to the other two species, reflecting the known species phylogeny.

**Fig. 7.**
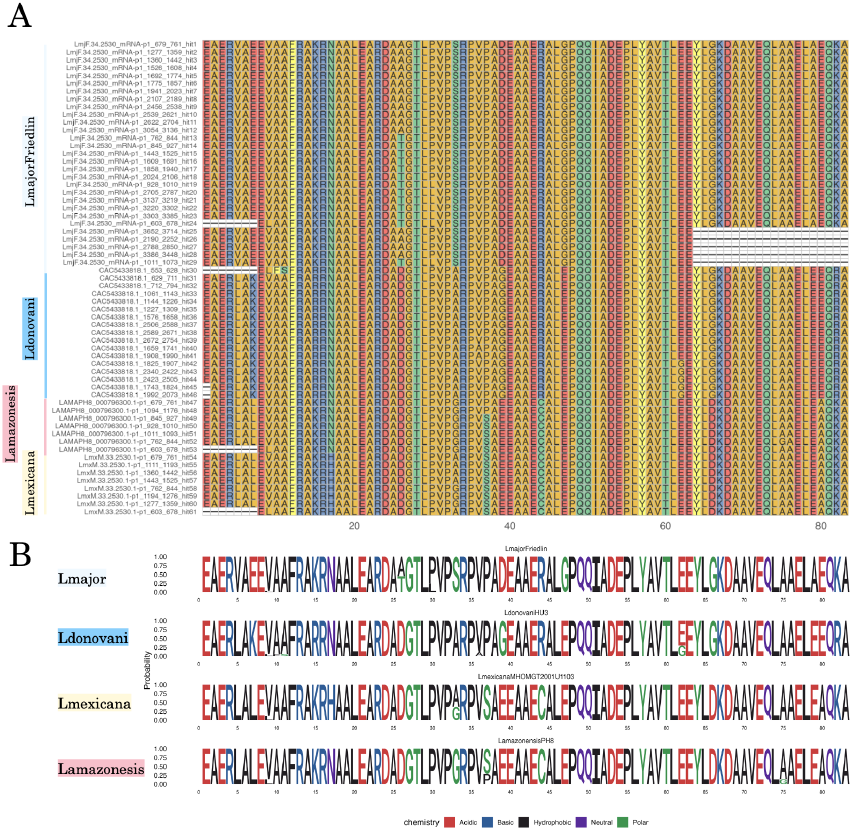
Comparative analysis of sequences in Orthogroup 1 containing LmjF.34.2530 (flagellar attachment zone protein). Multiple sequence alignment of repeat unit sequences (A) and species-specific sequence logos (B) are shown.

**Fig. 8.**
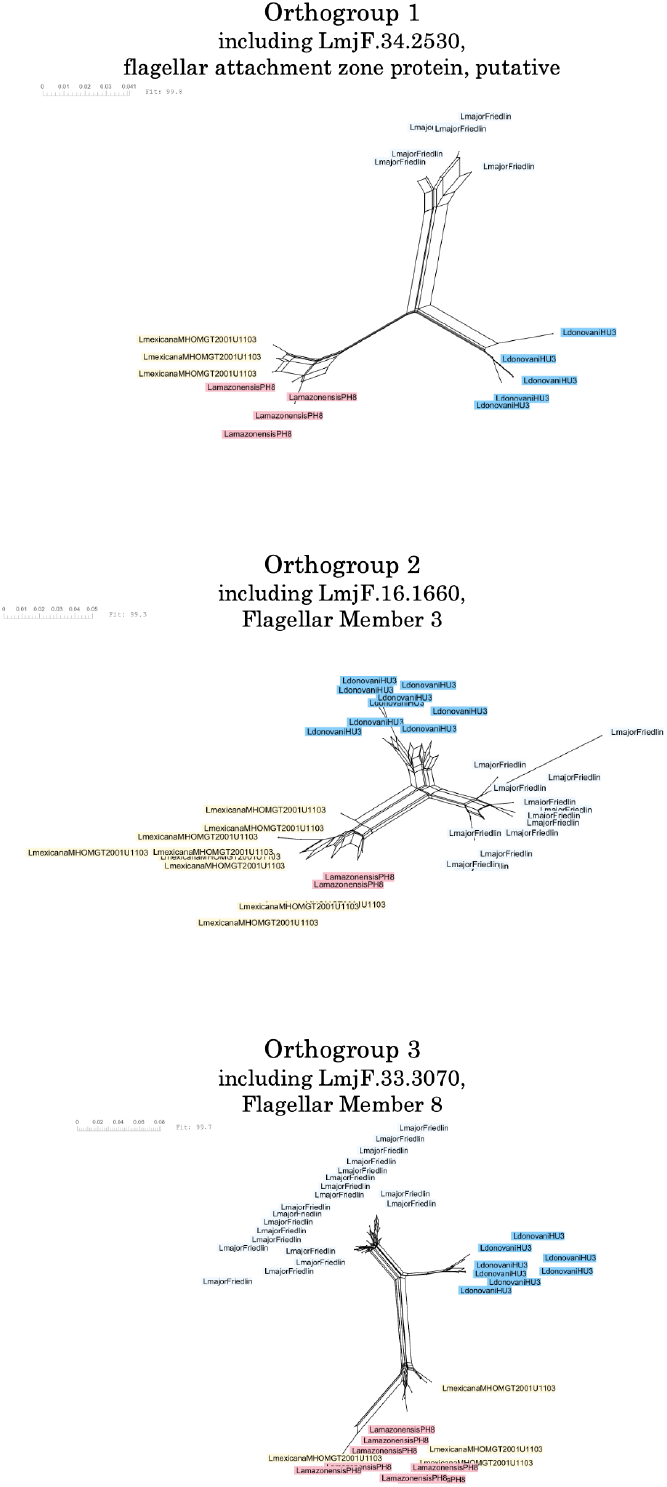
Phylogenetic networks of repeat unit sequences for the three orthogroups analyzed.

Analysis of repeat unit sequences revealed that amino acid differences were frequently shared across different repeat units within the same species (Fig. 7). Consistently, clustering of repeat unit sequences showed a clear separation by species rather than by repeat unit position or putative ancestral origin (Fig. 8).

## Discussion

In this study, we designed and quantitatively analyzed the repeat-related features of amino acid sequences from 12 species of *Plasmodium* and some kinetoplastids. Through these analyses, we identified a malaria-specific cluster (C1), as well as a *Trypanosoma*- and *Leishmania*-specific cluster (C2). Although the amino acid sequences in both clusters tended to contain large repeat regions, they exhibited contrasting trends in repeat-structure complexity; C1 showed high complexity whereas C2 exhibited significantly low complexity.

Functional analyses revealed that genes in C1 were enriched for AP2 domains, whereas those in C2 were enriched for flagellum-related GO terms and domains. In malaria parasites, the AP2 family constitutes one of the few classes of transcription factors and plays a critical role in gene expression regulation (24). Flagellar proteins have long been studied, primarily as motility-related components; however, recent studies have revealed that flagella are involved in a wide range of functions, including migration within host vectors, cell adhesion, and immune evasion (26, 27). In addition, C2 was enriched for DUF7623, a domain of unknown function. Therefore, our findings may provide useful insights into the functional roles of these uncharacterized domains.

Furthermore, C3 was enriched for the GO term host cell, with enrichment of genes encoding *Plasmodium*-exported proteins, including those associated with the PHIST family.PHIST proteins may contribute to host cell remodeling during the infection of human erythrocytes and are significantly expanded in *P. falciparum*, although their detailed functions remain poorly understood (28). Taken together, our results indicate that parasite-specific and functionally important proteins exhibit distinct repeat-related features, suggesting that lineage-specific evolution of repeat architectures may play a significant role in the acquisition of parasitic strategies in protozoa.

We further investigated why amino acid sequences in C2 maintain large repeat regions with low complexity by performing evolutionary analyses. One possible explanation for the low complexity is that these repeats were acquired relatively recently and have therefore accumulated only a limited number of mutations. However, comparisons of repeat sequences among orthologs from closely related species revealed species-specific amino acid substitutions, together with marked homogenization of repeat unit sequences within each species. If individual repeat units accumulated mutations independently, analogous to distinct gene sequences, repeat units derived from the same ancestral unit would be expected to show greater similarity across species than repeat units located at different positions within the same species. Under this scenario, repeat units would be expected to cluster according to their ancestral origin. However, our results do not support this expectation. Instead, amino acid differences were shared across different repeat units within the same species, and repeat unit sequences clustered primarily by species rather than by ancestral repeat units (Fig. 8). These patterns cannot be explained solely by the independent accumulation of mutations in individual repeat units and instead point to the presence of homogenizing forces acting on repeat unit sequences within each species. It is well established that repetitive sequences can undergo concerted evolution through mechanisms such as unequal crossing-over, leading to coordinated sequence evolution among repeat units (29, 30). Taken together, our results suggest that concerted evolution contributes to the maintenance of low-complexity repeat architectures in C2 proteins of *Leishmania*.

TRs composed of identical units (perfect repeats) tend to be less structured. In intracellular parasites, proteins containing perfect repeats are expressed at higher levels during those life-cycle stages which are capable of host cell invasion compared to other stages; these proteins are often associated with functions related to cell invasion (10). However, it has also been argued that perfect repeats do not necessarily imply specific structural or functional roles, but rather represent signatures of recent evolutionary events (6). The present results alone may be insufficient for determining whether the low complexity of repeat-structures is actively maintained by concerted evolution or its functional importance. Nevertheless, recent studies have shown that intrinsically disordered regions (IDRs) with low structural complexity can serve as scaffolds for molecular interactions. Future studies are required to elucidate the functional significance of these low-complexity repeat architectures in *Trypanosoma* and *Leishmania*.

Repeat architectures within protein sequences can exhibit highly complex patterns, arising from hierarchical repetition and diversification driven by mutational processes. Existing approaches are often insufficient to fully capture this structural complexity. To address this limitation, we developed a novel algorithm, Drepper, based on self dot plots and unsupervised learning, which enables quantitative characterization of repeat-structure complexity (Fig. 9). Although several tools have been proposed to analyze repeats using dot plots or related representations (21, 31–33), Drepper was specifically designed to extract repeat-related features that quantify architectural properties such as repeat-structure complexity. By providing a quantitative and integrative framework for analyzing complex repeat architectures, this approach sheds new light on the diversity and evolution of repeat-containing proteins and offers a foundation for future repeat analyses across diverse biological systems.

**Fig. 9.**
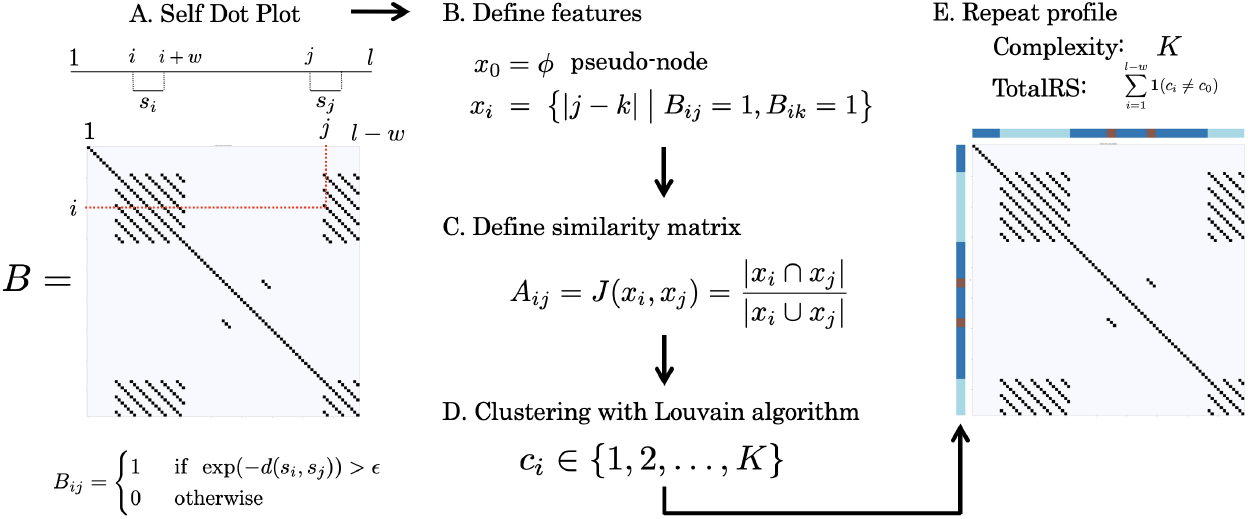
A schematic overview of the Drepper algorithm workflow.

## Conclusion

In this study, we analyzed repeat-related features, with a particular focus on repeat-structure complexity, using both existing tools and a newly developed algorithm, Drepper. We identified malaria-specific as well as *Trypanosoma*- and *Leishmania*-specific clusters of RPs. Genes within these clusters are likely to play important roles in parasite lineage, suggesting a close relationship between repeat architectures and parasite-specific strategies.

In the *Trypanosoma*- and *Leishmania*-specific cluster, repeat architectures exhibited low complexity; evolutionary analyses suggests that this pattern may be attributed to concerted evolution. Previous studies have discussed the role of repeat sequences in parasites in the context of host–parasite co-evolution (17), highlighting the potential impact of repeats on parasitism. In this study, we performed comprehensive repeat analyses using a novel algorithm, Drepper, from multiple perspectives. Drepper is freely available at https://github.com/hmatsu1226/Drepper.

Further studies are required to clarify the relationship between repeat architectures and parasitic strategies in protozoa. In this context, the application of advanced computational and bioinformatics approaches will become increasingly important. We anticipate that our study will serve as a foundation for future investigations in this field.

## Methods

### Overview

In this study, RPs were classified and subjected to downstream analyses based on quantitative features that characterize repeat architectures in amino acid sequences. To achieve this, it is essential to quantify repeat-related features from multiple perspectives. Accordingly, we developed a novel algorithm, Drepper, besides utilizing existing tools, to design and quantify diverse sets of repeat-related features. In the following sections, we first describe the features derived from the existing tool TANTAN, then introduce those computed using Drepper, and finally present the overall analytical workflow.

### Design of features based on TANTAN

Several sequence analysis tools have been developed to detect TR regions and their motif sequences in DNA and amino acid sequences (20). In this study, we employed TANTAN, which can handle amino acid sequences (22). TANTAN detects multiple TR regions within each amino acid sequence.

For a given amino acid sequence, suppose that *T* TRs are detected. For the *i*-th TR (*i*∈ [1, *T*]), TANTAN provides information on the start position *s*_*i*_, end position *e*_*i*_, repeat unit length *l*_*i*_, repeat copy number *r*_*i*_, motif sequence, and the actual sequences of individual repeat units. Based on the output of TANTAN, we designed the following TR-related features for each amino acid sequence:

**TRcount** The number of detected TR regions corresponding to *T*.

**MaxTRS** The maximum TR region size, defined as max_*i*_(*e*_*i*_ − *s*_*i*_). The index of the TR corresponding to MaxTRS is denoted by *i*^∗^ = arg max_*i*_(*e*_*i*_ − *s*_*i*_).

**RUP** A metric that quantifies repeat unit purity within the MaxTRS region. This value is computed based on the average normalized edit distance as follows:

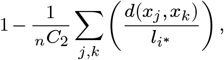

where 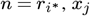 denotes the sequence of the *j*-th repeat unit of the *i*^∗^-th TR, and *d*(*a, b*) represents the edit distance between sequences *a* and *b*.

**MaxRUS** The maximum repeat unit size, defined as max_*i*_ *l*_*i*_.

### Design of features based on Drepper

We developed a novel algorithm, Drepper, based on self dot plots and unsupervised learning, which enables quantitative characterization of repeat-structure complexity (Fig. 9). Below, we describe how the proposed algorithm quantifies the repeat-structure complexity and related properties for an amino acid sequence *s* of length *l*. First, we defined the length-*w* substring from position *i* to *i* + *w* as *s*_*i*_ (*i*∈ [1, *l* −*w*]) and constructed a (*l* −*w*) × (*l* −*w*) distance matrix *D*, where each entry *D*_*i,j*_ is defined as the edit distance between sub-strings *s*_*i*_ and *s*_*j*_. We then defined a similarity matrix *S* by *S*_*i,j*_ = exp(− *D*_*i,j*_). Next, we binarized *S* to obtain a matrix *B* such that *B*_*i,j*_ = 1 if *S*_*i,j*_ > *ϵ*, otherwise *B*_*i,j*_ = 0 (Fig. 9A). In this study, we set *w* = 10 and *ϵ* = 0.2.

Next, from the matrix *B*, we defined for each position *i* ∈ [1, *l* −*w*], a feature vector *x*_*i*_, as follows (Fig. 9B):

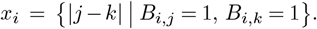

This is the set of pairwise distances among all indices *j* for which *B*_*i,j*_ = 1 is true. If a given substring *s*_*i*_ has a very large number of similar substrings (i.e., the number of indices *j* satisfying *B*_*i,j*_ = 1 is very large), computing all pairwise distances becomes computationally expensive. In such cases, we restrict the construction of *x*_*i*_ to a subset of indices *j* corresponding to the top 50 smallest values of | *i* − *j* |.

We then added a pseudo-data point representing unique, non-repetitive sub-sequences by introducing *x*_0_ = *ϕ*, where *ϕ* denotes an empty set. We defined a similarity matrix *A* among the (*l* − *w* + 1) feature vectors using the Jaccard index (Fig. 9C):

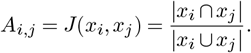

We defined *J*(*ϕ, ϕ*) = 1. Subsequently, we binarized the similarity matrix *A* at a threshold of 0.5 to obtain *A*^*′*^. Next, *A*^*′*^ was treated as the adjacency matrix of a graph, and clustering was performed using the Louvain algorithm. This procedure yielded cluster assignments *c*_*i*_ for each node (*i* ∈ [0, *l* − *w*]) (Fig. 9D). Here, *c*_1_, …, *c*_*l*−*w*_ represent the clustering results for each position in the self dot plot, whereas *c*_0_ corresponds to the cluster ID, representing regions that do not correspond to repeats.

As described above, positions within a sequence can be clustered based on information from the off-diagonal elements of the self dot plot, enabling the quantitative characterization of properties such as the complexity of repeat structures in the amino acid sequence. Based on this framework, we designed the following features to quantify repeat-related features (Fig. 9E).

**Complexity** The complexity of repeat regions within an amino acid sequence, quantified as the diversity of clusters, is defined as: *K* = max_*i*_(*c*_*i*_).

**TotalRS** The total extent of repeat-associated regions within an amino acid sequence is computed as: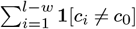

### Datasets and pre-processing

We analyzed the amino acid sequences from 12 species of malaria parasites and some kinetoplastids, focusing on strains with relatively well-curated genome annotations. The target species and their corresponding strains are listed in Table 1. Amino acid sequence data for malaria parasites were obtained from PlasmoDB (34) (release 68), and those for kinetoplastids were obtained from TriTrypDB (35) (release 68). Amino acid sequences encoded by short contigs, mitochondrial genomes, or apicoplast chromosomes were excluded from the analysis.

For each of the 12 species, the repeat-related features were computed for each amino acid sequence using both TANTAN and Drepper. Together with the sequence length (Length), seven features were calculated for each amino acid sequence. Features from all species were integrated to construct a data matrix *X* of size *N×* 7 for *N* amino acid sequences. All features except RUP were log-transformed. In addition, amino acid sequences with TotalRS equal to zero were removed prior to downstream analyses. The final dataset comprised *N* = 6,380 sequences of RPs.

Using this feature matrix, amino acid sequences were clustered using the *k*-means algorithm, and principal component analysis (PCA) was performed on *X*. To enable fine-grained characterization of RP properties based on repeat-related features, we set the number of clusters to a relatively large value, *K* = 7.

### Enrichment analyses

#### Species enrichment analysis

Let *C* denote a 12 × 7 count matrix summarizing the number of RPs from each of the 12 species assigned to each cluster. We tested whether a specific group of species was enriched in a given cluster, using the following procedure.

Let *g* be a vector representing species group membership, where *g*_*i*_ = *A* if species *i* belongs to the focal species group, otherwise *g*_*i*_ = *B*. Let *k* denote the cluster of interest; define a vector *x* such that *x*_*i*_ = *C*_*i,k*_ represents the number of RPs from species *i* assigned to the cluster *k*. The total number of RPs for species *i* is defined as 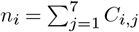.

We tested whether species with *g*_*i*_ = *A* exhibited a significantly higher proportion of RPs assigned to cluster *k*, i.e., *x*_*i*_*/n*_*i*_, compared to species with *g*_*i*_ = *B*. Statistical testing was performed using a generalized linear model based on a binomial distribution implemented in R (`glm()`). The response variable was specified as `cbind` (*x*_*i*_, *n*_*i*_ −*x*_*i*_), the explanatory variable was *g*_*i*_, and the model was fitted with `family = binomial`. The *p*-value was obtained from the significance of the regression coefficient associated with *g*_*i*_.

#### GO enrichment analysis

GO annotations for each species were obtained from GAF files downloaded from PlasmoDB and TriTrypDB. We focused on 1,963 GO terms that appeared at least once among the 6,380 RPs analyzed.

For each of the 1,963 GO terms, we counted the number of RPs associated with that term in each cluster. The enrichment of a given GO term within a specific cluster was assessed using a one-sided Fisher’s exact test. For each cluster, raw *p*-values were computed for all GO terms and subsequently adjusted for multiple testing using Bonferroni correction.

Two control sets were used for comparison: (1) the set of amino acid sequences belonging to all clusters other than the target cluster and (2) the set of all amino acid sequences not belonging to the target cluster, including non-RPs, that appeared at least once in the GAF files. The enrichment *p*-values were computed separately, using each control set.

#### Protein signature enrichment analysis

Protein signatures present in each amino acid sequence were predicted using InterProScan (release 5.76–106.0) (36). We focused on 6,442 signature accessions that appeared at least once among the 6,380 RPs.

For each signature accession, we counted the number of RPs associated with that accession, in each cluster. When the same signature accession was predicted in multiple regions of a single amino acid sequence, it was counted only once for that sequence. The enrichment of a given signature accession within a specific cluster was assessed using a one-sided Fisher’s exact test, followed by Bonferroni correction to obtain adjusted *p*-values.

Two control sets were used: (1) the set of amino acid sequences belonging to all clusters other than the target cluster and (2) the set of all amino acid sequences not belonging to the target cluster, including non-RPs, for which at least one protein signature was predicted by InterProScan. The enrichment *p*-values were computed separately using each control set.

### Evolutionary analysis

Evolutionary analyses were restricted to nine kinetoplastid species. Orthologous relationships were inferred using OrthoFinder (37) and orthogroups that were frequently assigned to C2 were extracted. Among the amino acid sequences belonging to the same orthogroup, all repeat motifs predicted by TANTAN in the *Leishmania major* Friedlin strain were treated as candidate repeat motifs. For each candidate motif, all amino acid sequences in the same orthogroup were used as references, and the motif sequence was used as a query in BLASTP searches to enumerate all sub-sequences matching the motif. An e-value threshold of 1 × 10^−5^ was applied, followed by the filtering of hits with bit scores below 100. The resulting sequences were output in FASTA format.

Similar sequences for each motif were listed, and those motifs for which similar sequences were detected in all four *Leishmania* species (LmajorFriedlin, LdonovaniHU3, LamazonensisPH8, and LmexicanaMHOMGT2001U1103) were selected; the corresponding hit sequences were subjected to comparative sequence and phylogenetic network analyses. Multiple sequence alignment was performed using MAFFT (38). The resulting alignments were visualized globally using ggmsa (39) and sequence logos were generated for each species using ggseqlogo (40). Identical sequences within the same species were collapsed into a single representative sequence, and sequence similarity was visualized as a phylogenetic network using SplitsTree (41).

## ACKNOWLEDGEMENTS

This work was supported by the Japan Society for the Promotion of Science (grant number 21K17858 to Hirotaka Matsumoto and 25K18791 to Jing Hong).

## Supplementary Note 1: Supplementary information for clustering of RPs

The density distributions of the 12 species in the PCA space are shown in Fig. 10. In *Plasmodium*, particularly *P. falciparum*, a relatively large number of RPs are located in the regions corresponding to C1. In contrast, in *Trypanosoma* species (TbruceiTREU927 and TcruziBrazilA4) and multiple *Leishmania* species (LmajorFriedlin, LdonovaniHU3, LamazonensisPH8, and LmexicanaMHOMGT2001U1103), a relatively large number of RPs are distributed in regions corresponding to C2.

**Fig. 10.**
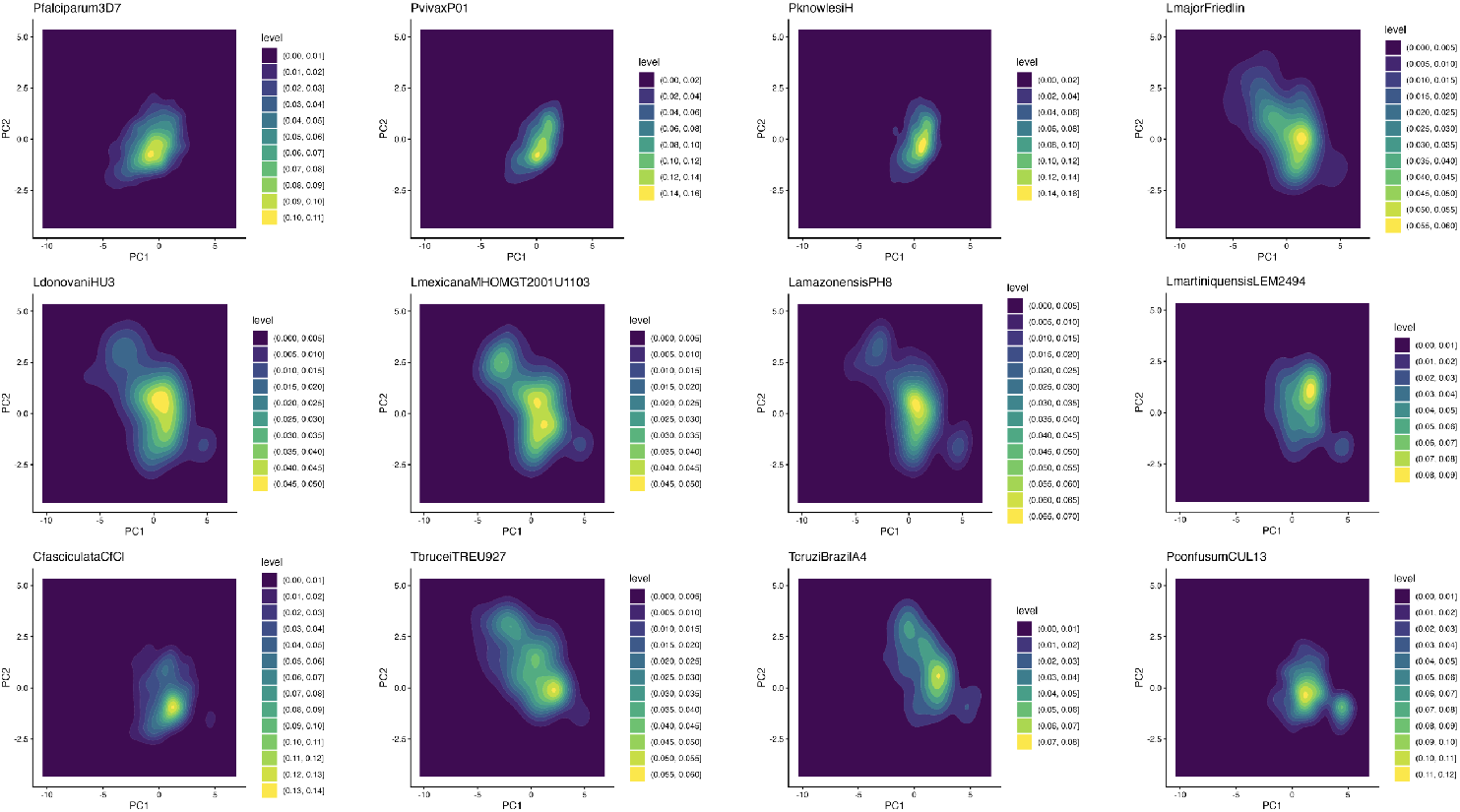
Density distributions of RPs in the PCA space, shown for all species.

The density distributions illustrating the relationship between TotalRS and Complexity across all 12 species and clusters are shown in Figs. 11 and 12, respectively. In *Plasmodium*, the complexity tends to increase with increasing TotalRS. By contrast, in *Trypanosoma* species (TbruceiTREU927 and TcruziBrazilA4) and multiple *Leishmania* species (LmajorFriedlin, LdonovaniHU3, LamazonensisPH8, and LmexicanaMHOMGT2001U1103), a substantial number of RPs exhibit large TotalRS values but low complexity. As described in the main text, C1 and C2 both consist of RPs with large TotalRS values, but differ markedly in complexity, with C1 showing high and C2 showing low complexity. C3 and C4 show intermediate TotalRS values, with C3 exhibiting slightly higher complexity than C4. C5 and C6 generally show low values for both TotalRS and complexity. Representative examples of RPs from each cluster, visualized using dot plots, are shown in Fig. 13. RPs in C3 are characterized by moderately large TotalRS values and relatively high complexity. RPs in C5 and C6 generally have short amino acid sequence lengths. In C7, the repetitive sequences are distributed in a dispersed manner rather than forming contiguous tandem repeat regions.

**Fig. 11.**
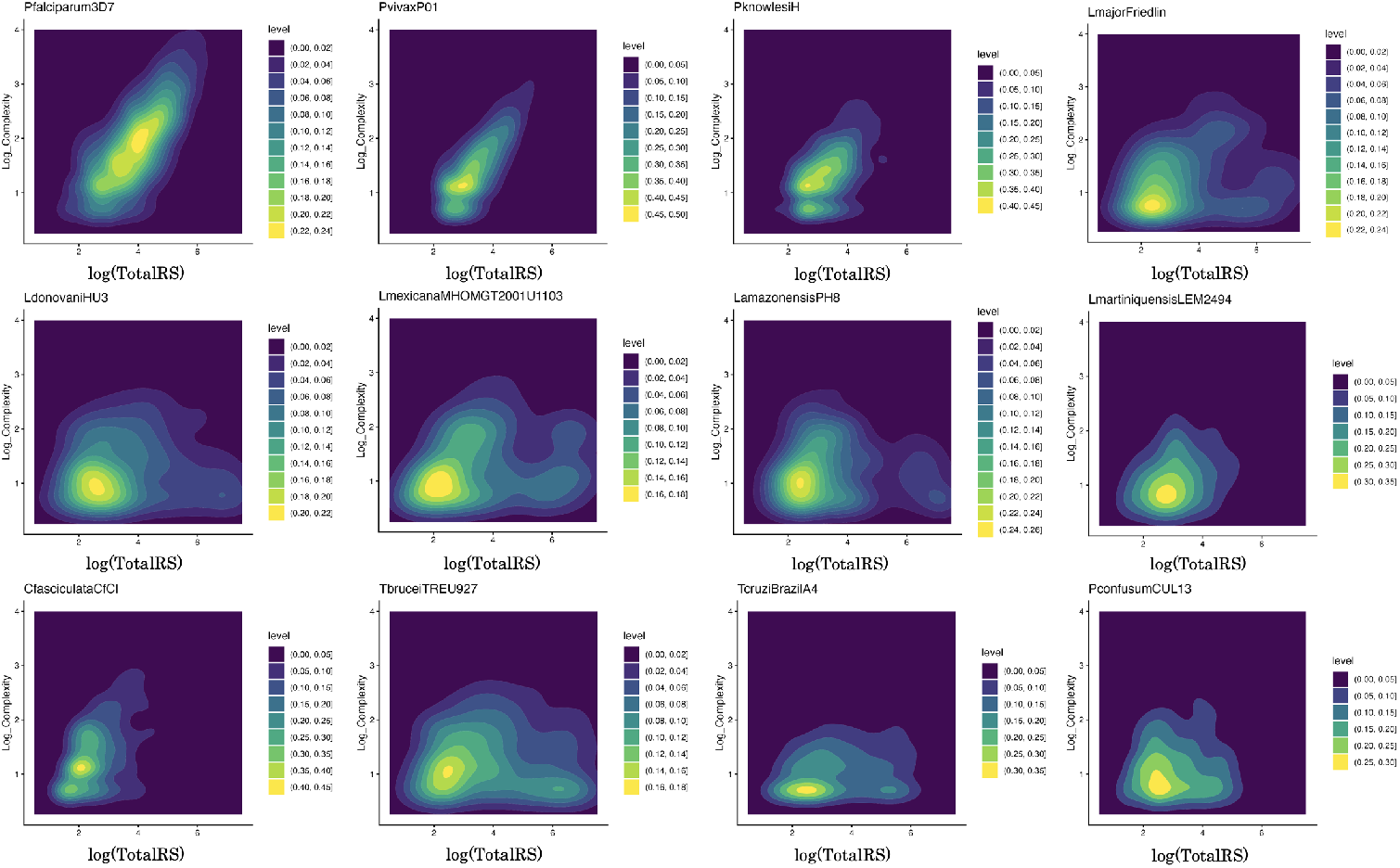
Density plots of RPs for all species, with the logarithm of TotalRS on x-axis and logarithm of Complexity on y-axis.

**Fig. 12.**
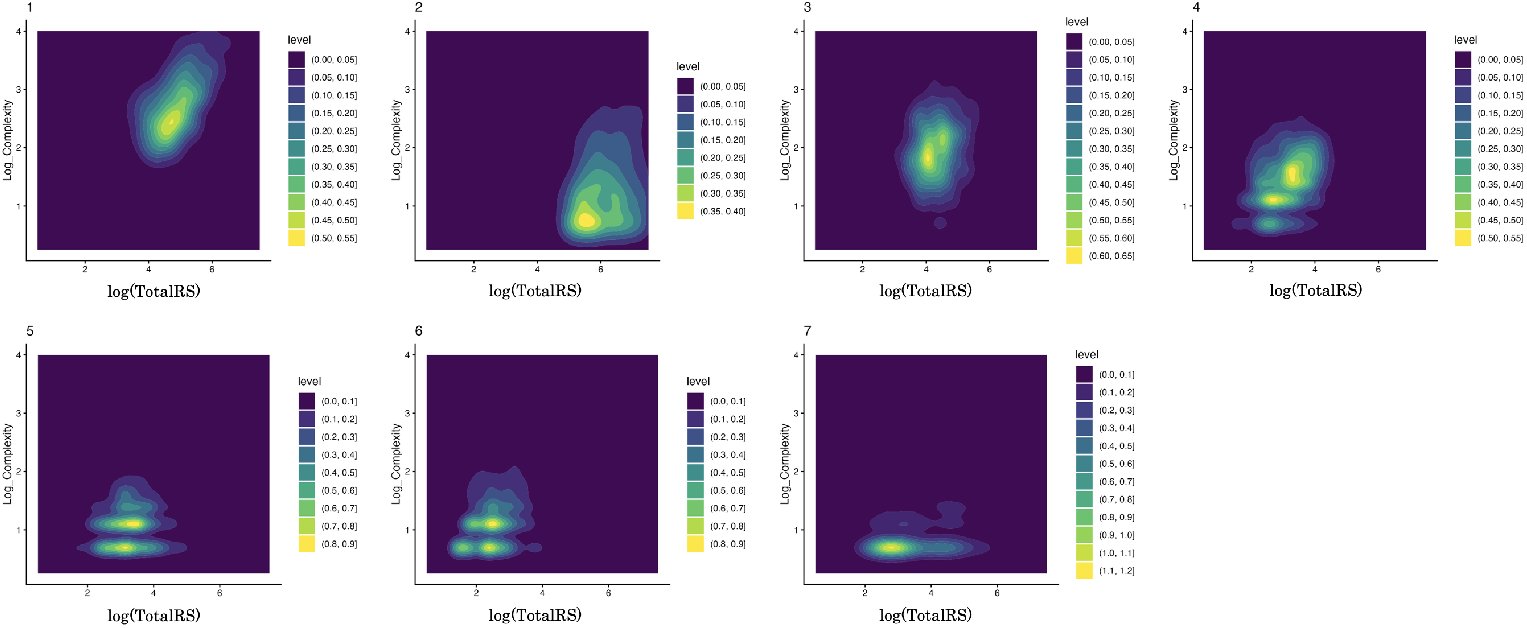
Density plots of RPs for all clusters, with the logarithm of TotalRS on x-axis and logarithm of Complexity on y-axis.

**Fig. 13.**
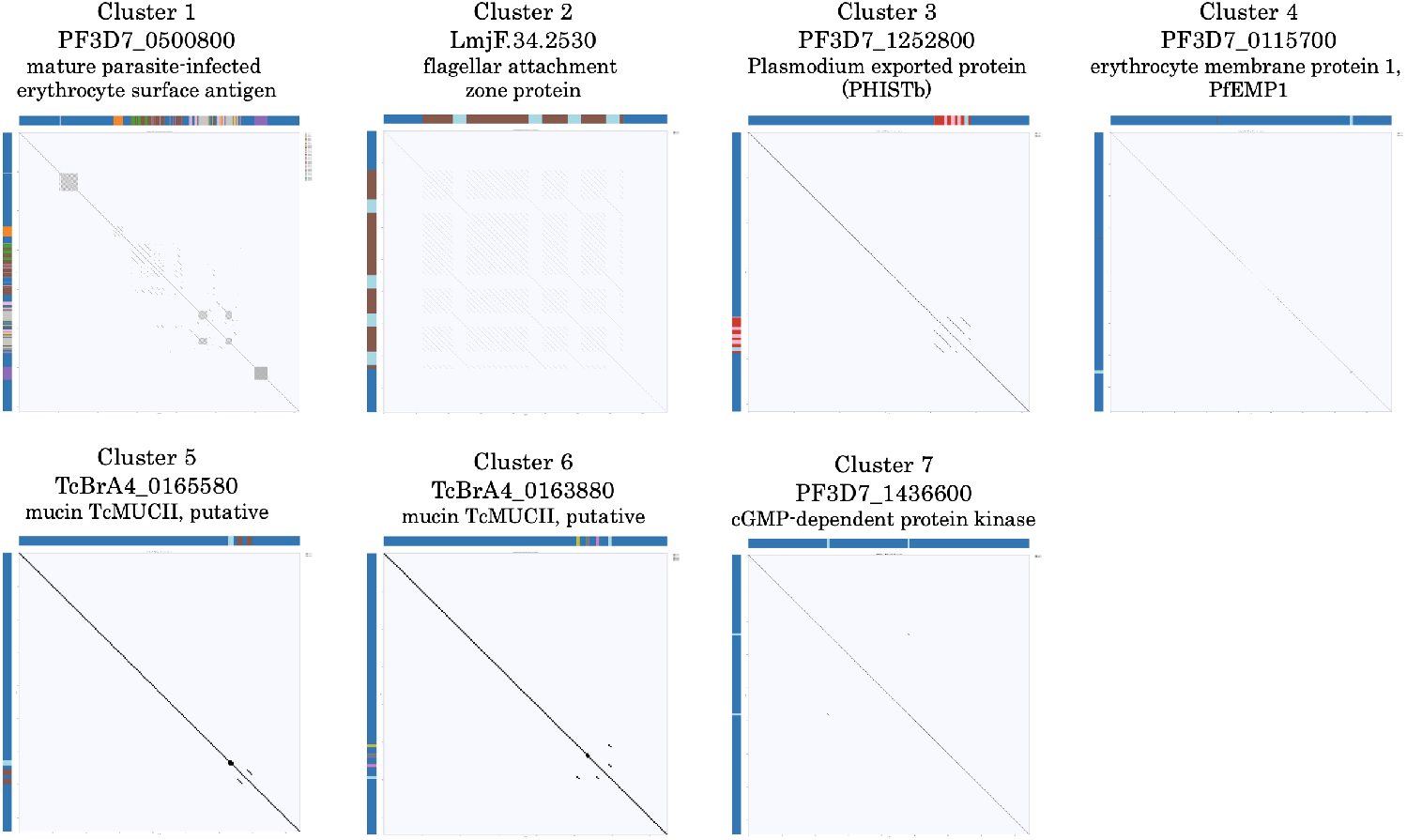
Representative dot plots of RPs in all clusters. The annotations displayed along the top and left margins of the dot plot, together with their colors, represent the clustering results obtained by Drepper (see Methods section).

## Supplementary Note 2: Supplementary information for evolutionary analysis

Multiple sequence alignments and species-specific sequence logos for Orthogroups 2 and 3, as discussed in the main text, are shown in Fig. 14. In both orthogroups, multiple positions exhibit species-specific homogenization of amino acid substitutions. For example, in Orthogroup 2, the amino acid at position 11 is homogenized as alanine (A) in LmajorFriedlin and LdonovaniHU3, whereas it is homogenized as threonine (T) in LmexicanaMHOMGT2001U1103 and LamazonensisPH8.

**Fig. 14.**
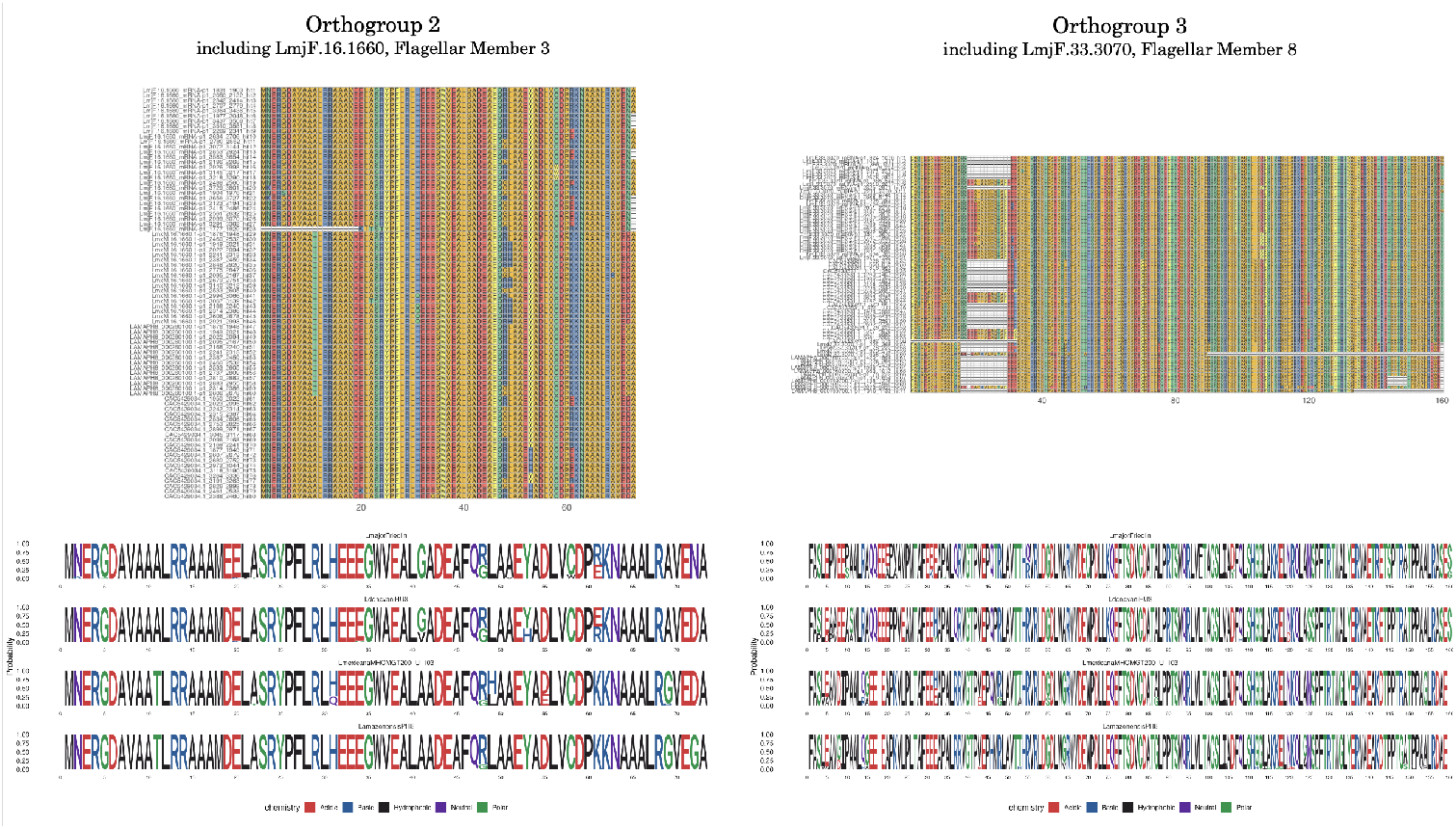
Comparative analysis of sequences in Orthogroup 2 (left) and Orthogroup 3 (right). Multiple sequence alignment of repeat unit sequences (top) and species-specific sequence logos (bottom) are shown.

